# Processing at Phrase Boundaries During Self-Paced Reading

**DOI:** 10.64898/2026.07.13.738177

**Authors:** Jenna Hooper, Julia Dengler, Nicoletta Biondo, Jacob Odeh, Pooja Mudigonda, Rajesh Kana, David Basilico, Matthew John Nelson

## Abstract

Studies of the difficulty of Object-Relative sentences (ORs) have yielded important insights about the nature of sentence parsing. Given the incremental nature of sentence parsing, understanding the precise locations of OR processing difficulty is a critical aspect of this. Prior studies have frequently highlighted the Relative Clause (RC) verb as a major locus for the OR penalty. However, it is difficult to disentangle gap-filing and phrase wrap-up effects from the presence of a verb, which all coincide in typical ORs. Here we introduce prepositional verbs like “spoke to” in OR sentences in a one word-at-a-time self-paced reading experiment to disentangle this. The OR penalty primarily occurred at the phrase-ending preposition rather than the verb, with some of the penalty also occurring at the verb, reflecting that gap-filling and phrase wrap-up underlie more of the effect than processes occurring at the verb itself. However, repeated exposure to these sentences over the course of a session led to an increasing fraction of the OR penalty moving to the verb before the preposition, likely resulting from local-experience-driven prediction. Further repeated exposure eventually diminished, but did not eliminate, the total combined penalty. This highlights that the location of processing difficulty is itself plastic and malleable with experience. We further found that semantic congruence most strongly affected RTs at these key locations of OR sentences compared to SR and control sentences, suggesting that semantic processes contribute to the OR penalty. More syntactic nodes being closed at the phrase boundary also lead to increased phrase boundary RTs for all RC sentences, suggesting that the amount of material being integrated at RC boundaries also drives the RT cost there. Together, these findings advance our understanding of the processes underlying the OR penalty.

## Introduction

Linguistic theories of sentence processing propose that sentences are read as hierarchical structures of nested phrases rather than a linear sequence of words (Berwick et al., 2013; Chomsky, 1957; Haegeman, 2005; Sportiche et al., 2013). In this theory, constituents, or groups of words that function as a unit, are built and completed while reading. The places where the constituents are completed, or phrase boundaries, are proposed to be important locations of integration, where both semantic and syntactic information are processed. However, phrases and phrase boundaries investigated through self-paced reading have been underexplored in the literature.

In investigating behavioral results of periodic neural activity of self-paced reading, Lo et al., (2023) found that that participants’ response times slow at the phrase boundaries. This is likely due to chunking, where the participant pauses to consolidate information together to begin new chunking. This process can decrease memory demands, leading to easier processing of a sentence.

Neurophysiological evidence also supports the view of comprehension as an incrementally built phrase structure. Nelson et al., (2017) demonstrates a neural merging process where the parsing of constituents occurs, occurring shortly after the last word of each syntactic constituent. These results support that there is a location at phrase boundaries where integration occurs, making these boundaries good targets for further studying of sentence processing mechanisms.

In psycholinguistics, differences in processing difficulty between subject relative clauses and object relative clauses have been studied to understand processing of complex syntactic comprehension.

1. The cat that chased the dog was brown.
2. The cat that the dog chased was brown.

These sentences demonstrate two kinds of relative clauses. In subject relative sentences (Sentence 1), the head noun phrase (NP) “cat” is the subject of the relative clause “that chased the dog”. In object relative sentences (Sentence 2), the head NP “cat” is the object of the relative clause “that the dog chased”.

There is a large ongoing discussion of why object relative sentences are more difficult to process than subject relative sentences, and this asymmetry has been found consistently across various experimental methods (Caplan & Waters, 1998; Caramazza & Zurif, 1976; King & Just, 1991). Several theories have been proposed to explain this phenomenon, and they fit into three categories: frequency-based, memory-based, and semantic accounts (Gordon & Lowder, 2012). Frequency-based accounts describe that the frequency of syntactic structures impacts their difficulty, with less encountered structures being more difficult to process. Memory-based accounts describe difficulty arising from encoding and retrieving syntactic information during comprehension. Semantic based accounts describe that the meaning of the grammatical components can influence the ease of understanding the sentence.

### Frequency

One explanation for the higher difficulty in processing ORCs focuses on differences in experience with different sentence structures. Frequency-based accounts describe that readers find it easier to process sentence structures they have encountered most often. Corpus analysis supports this view by showing that that SRCs occur more frequently in English than ORCs (Gordon & Hendrick, 2005; Roland et al., 2007). Due to more experience processing SRCs, readers may find those structures easier. Frequency-based accounts are further supported by Reali & Christiansen (2007)’s finding that processing difficulty does not just depend upon the sentence structure, but also the frequency of types of pronouns within noun phrases. In English, ORCs are more likely than SRCs to contain a personal pronoun as the embedded noun phrase.

1. [SRC] “The reporter that attacked you admitted the error.”
2. [ORC] “The reporter that you attacked admitted the error.”

Reali & Christiansen (2007) found that when SRC and ORC sample sentences use personal pronouns (see Sentences (1) and (2)), participants read ORCs faster than SRCs. This result demonstrates that ORC difficulty cannot be fully explained by memory constraints alone. Instead, it suggests that frequency of sentence constructions and one’s experience with them can impact comprehension speed.

This account is further supported by Wells et al., (2009)’s findings that participants who practiced reading ORC later processed these structures more quickly, which suggest that experience can reduce comprehension difficulty. Similarly, MacDonald & Christiansen, (2002) found that that the highly frequent N-V-N sequence present in SRCs may contribute to their ease, compared to the less common N-N-V-V of ORCs. Other frequency-based accounts elaborate that the larger frequency of SRC structures leads to greater certainty reading them and thus making them easier to predict than ORCs.

There are some troubles with this frequency-based explanation, as it does not fully explain the processing asymmetry between ORCs and SRCs. Gordon & Hendrick (2005) note how in corpus data, SRCs often have an intransitive embedded verb, while ORCs always have a transitive one. When SRC with an intransitive verb are removed from the corpus, the difference in frequency between SRC and ORC decreases greatly. This raises the “grain problem” (Gordon et al., 2004; Mitchell et al., 1995), which argues that frequency-based theories should specify how experience is influencing processing.

### Memory

Memory-based accounts argue that the ORC-SRC asymmetry is associated with memory limitations during complex sentence processing, describing greater cognitive demands on memory. Unlike SRCs, ORC head nouns cannot be immediately integrated into the sentence, and it instead must be held in working memory until encountering the embedded verb and establishing grammatical roles. This additional process is considered to make sentence processing more difficult.

The early theories concerning memory-based accounts mainly attributed the asymmetry to limitations on working-memory capacity. These accounts argued that readers must devote cognitive resources to maintaining unattached noun phrases while simultaneously processing the rest of the sentence, increasing the possibility that memory capacity will be overloaded (Just & Carpenter, 1992; King & Just, 1991).

More recent studies have broadened from just working memory, and have focused on linguistic influences on memory retrieval processes. The cue-based parsing framework (Lewis et al., 2006; Lewis & Vasishth, 2005; Van Dyke & Lewis, 2003) describes reader using linguistic cues at the embedded verb to retrieve noun phrases from memory and solve a long distance dependency. ORCs are particularly difficult because the parser must retrieve the correct noun phrase from among competing noun phrases. Retrieval becomes more difficult when the noun phrases exhibit similar characteristics and are thus harder to distinguish.

Two major theories arose to explain the impact of memory demands on processing difficult of ORCs. One is the dependency locality theory (Gibson, 1998, 2000; Grodner & Gibson, 2005) which describes comprehension becoming more difficult as the number of intervening discourse referents increases between the NP and its integration with a verb. In object relative clauses, the head noun must remain active across multiple intervening discourse referents before it can be integrated with the embedded verb. Sentences with more intervening referents require greater memory resources and are processed less efficiently, thus explaining a cause for the greater ORC processing difficulty. The dependency locality theory is supported by the givenness hierarchy in which high accessible referents, like “I” or “you”, have relatively low processing costs due to their established existence in the discussion. In contrast, less accessible referents, like specific noun phrases like “the senator”, have higher processing costs due to the effort of maintenance and retrieving, and thus increases memory demands.

A second theory is the similarity-based interference account, which argues that processing difficulty depends less on distance between noun phrases and instead on their similarity (Gordon et al., 2001, 2002, 2004; Van Dyke & Lewis, 2003; Van Dyke & McElree, 2006). While reading, both noun phrases are stored in memory until encountering the embedded verb, where the reader would then assign the appropriate thematic roles, such as agent and patient roles. When the noun phrases are very semantically or syntactically similar, they are easier to confuse and their thematic role assignment becomes more difficult. Therefore, ORC processing difficulty is described to arise from retrieval problems, and not just memory storage constraints.

### Semantics

Semantic accounts argue that processing difficulty is influenced by the meaning relationships in a sentence, in which the meaning of a sentence is more straightforward or natural with a SRC than an ORC. In these theories, comprehension is easier if the semantic properties support the expected syntactic interpretations, and harder if the semantic relationships are less common or likely.

One line of research suggests that the SRC versus ORC asymmetry is decreased when there is an inherent semantic relationship between the critical noun phrases. King & Just (1991) found that readers experienced less difficulty processing object relative clauses when the nouns shared semantic features, compared with sentences in which the relationship between the nouns was random. These results show that semantic information can impact sentence interpretation and somewhat decrease the processing difficulty typically experienced with ORCs.

A related explanation describes a perspective shift during comprehension. MacWhinney & Pléh, (1988) proposed that object relative clauses modifying a subject head noun require readers to shift perspective while constructing the sentence representation. This additional perspective shift is not present in subject relative clauses, making SRCs comparatively easier to understand.

Another semantic approach focuses on animacy of the critical noun phrases in relative clause processing (Traxler et al., 2002). The general results show that ORC is particularly difficult when the unattached head noun is animate and the embedded noun is inanimate. However, this processing asymmetry is decreased when the head noun is inanimate and the embedded noun is animate, which suggests that readers’ expectations about the plausibility of thematic roles also influence comprehension.

Traxler et al., (2002) proposed these findings in relation to the active filler strategy, which describes that readers initially predict a relative clause will follow the more common subject relative clause structure when encountering the complementizer “that”. When this expectation is incorrect, the sentence must be reanalyzed. Reanalysis is more difficult when the extracted head noun is animate because animate nouns are often interpreted as agents rather than as patients. In contrast, inanimate noun phrases are more easily interpreted as patients or objects, which makes reanalysis less difficult.

### Phrase boundaries

ORCs are shown to be more difficult than SRC and control sentences throughout many studies, but none investigate the effect of the phrase boundary for ORCs and SRCs. Their processing difficulty polarity is known, but exactly why is still not fully understood, which makes them interesting stimuli for comparison of syntax and sentence comprehension.

Psycholinguistic theories generally assume that sentence comprehension proceeds through the incremental construction of hierarchical constituent structure, rather than by processing words in isolation (Frazier & Fodor, 1978; Frazier, 1987; Gibson, 1998; Phillips, 2003; Pritchett, 1992). Under this view, phrase boundaries are natural points at which syntactic structure is completed and integrated with the developing sentence representation.

Despite the central role of constituent structure in theories of parsing, most self-paced reading studies have localized processing difficulty with respect to lexical regions (e.g., the verb or a critical noun) rather than equivalent structural positions across sentence types. Previous studies comparing ORCs and SRCs have typically interpreted processing difficulty with respect to lexical regions, particularly the embedded verb, where retrieval and integration operations are assumed to occur.

In the present study, we focus on the boundary marking completion of the subject noun phrase. For example, in the subject-relative sentence *The cat that chased the dog | was brown*, the boundary follows the embedded relative clause, whereas in the object-relative sentence *The cat that the dog chased | was brown*, the boundary occurs at the same structural position despite the different word order. Likewise, in a canonical sentence such as *The brown cat | chased the dog*, the corresponding boundary marks completion of the subject noun phrase. Thus, although these sentence types differ lexically and syntactically, they share a common structural boundary that can be used as a point of comparison.

Aligning reading times to this shared structural position allows processing to be compared at equivalent stages of syntactic structure building across sentence types, rather than at corresponding word positions or lexical categories. If processing difficulty is tied primarily to constituent completion, reading-time effects should remain aligned to this boundary even when the lexical realization of the boundary differs.

To test this prediction, we included object-relative sentences in which the constituent boundary coincided either with the main verb or, for prepositional verbs, with the following preposition. This manipulation dissociates structural position from lexical category, allowing us to determine whether the locus of processing difficulty is better characterized as a property of constituent completion than of individual lexical items.

Therefore, should the object relative clause difficulty be associated with processes like memory retrieval and semantic interpretation, these effects should be observable at the phrase boundaries. To study the effect of the phrase boundary for ORCs and SRCs, we created a self-paced reading task to study response times and processing of ORCs, SRCs, and control sentences.

## Methods

### Participants

We collected self-paced reading data from 35 healthy, native English-speaking participants aged 18–25 years (M = 19.5, SD = 1.7). Participants were undergraduate students, with the exception of two participants who held bachelor’s degrees and were enrolled in postgraduate programs. Based on self-reported gender, the sample included 22 women, 12 men, and 1 nonbinary participant. Of the participants, 30 reported being right-handed and five reported being left-handed; two of the right-handed participants also reported some degree of ambidexterity. All study procedures were approved by the Institutional Review Board at the University of Alabama at Birmingham (IRB protocol no. 300004963; approved April 13, 2023). Participants reviewed an IRB-approved information sheet and provided informed consent in accordance with the approved protocol. Participants received $5 in compensation for completing the approximately 30-minute experiment.

### Experimental Procedure

We administered the experiment in EPrime 3.0 (Psychology Software Tools) on a Windows laptop with a touch screen. Participants were seated comfortably at a table with the laptop at a comfortable distance in front of them. We asked participants to read and comprehend sentences presented visually, one word at a time. Words were presented in the center of the screen in 100 point bolded Arial font with the first word of each sentence capitalized and the rest of the text in lowercase. Before each sentence, a plus sign was shown on the screen waiting for the participant to initiate the sentence. Participants pressed the space bar to advance past the plus sign and then to each subsequent word in the sentence. The experimenter explicitly instructed each participant to advance through the sentences at whatever pace they preferred, so long as they could successfully perform the task (i.e. select the correct picture after each sentence). We did not inform participants that their reading times would be analyzed. At the end of each sentence, participants were presented with four pictures, and their task was to choose which of the pictures corresponded to the sentence they just read. Participants responded by touching that picture on the laptop screen with their finger and their touch position was recorded. Participants completed 2 blocks of 80 sentences each, with a self-timed break between the 2 blocks.

### Sentence stimuli

The sentence stimuli included four sentence structures: object relative (OR) sentences (e.g., *The cat that the dog chased was brown*), subject relative (SR) sentences (*The cat that chased the dog was brown*), subject-modifier canonical (SMC) sentences (*The brown cat chased the dog*) and object-modifier canonical (OMC) sentences (*The cat chased the brown dog*). The content of the sentences was always imageable in the pictures presented at the end of the sentence. The 4 pictures presented simultaneously probed patient/agent assignment (i.e., who chased whom?) and noun-complement assignment (was the cat or the dog brown?) independently in a 2×2 design (Figure 1). In each trial 1 of the 4 pictures was correct, 1 picture reflected a agent-patient error, 1 reflected a noun-complement error, and 1 reflected both errors simultaneously. When the participant makes an error this allows us to understand which of these mappings were misunderstood.

**Figure 1.**
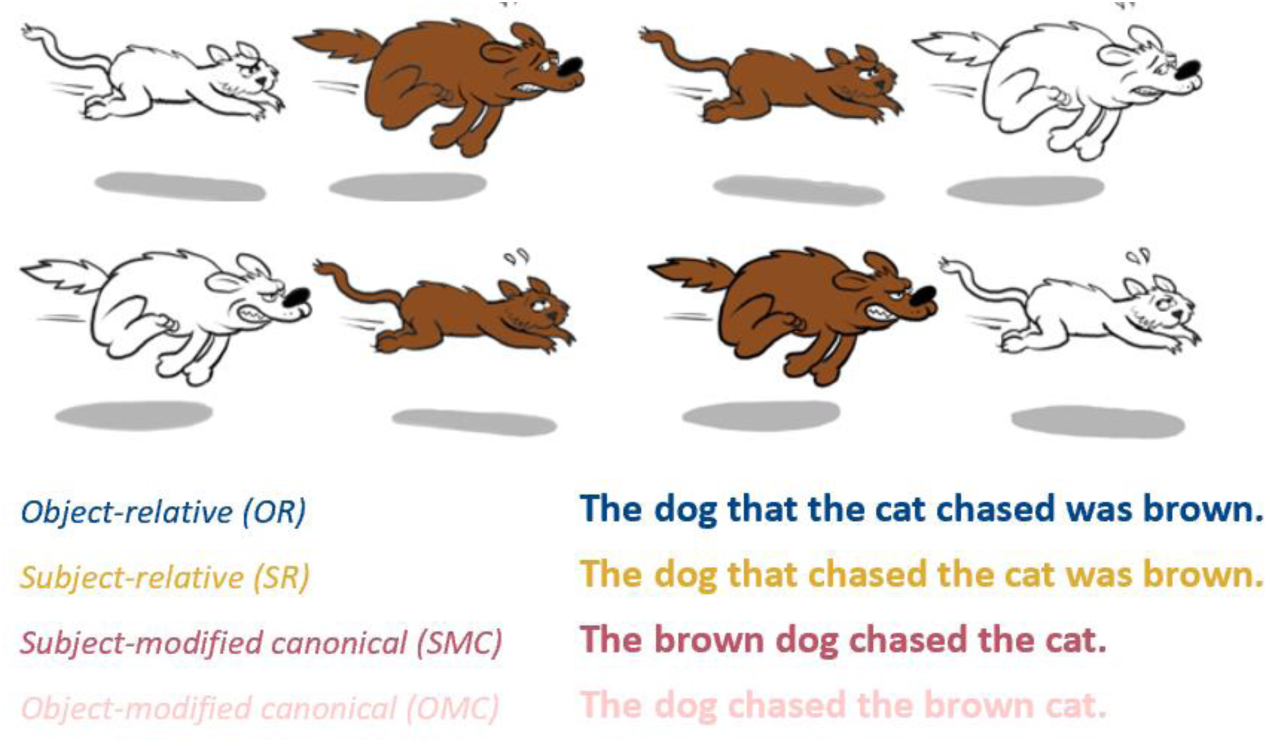
The four pictures show an example of the pictures shown after one of the sentences. The pictures comprise a 2×2 design independently probing the participant’s comprehension of the patient-agent relationship in the sentence (*Who is chasing whom?*) and the complement-noun relationship (*Is the cat or the dog brown?*). The sentences show the four different sentence structures presented.

Each sentence requires four semantic content roles-the first noun, the second noun, the verb and the noun-complement. All nouns in these content roles were animate and always introduced in the sentences with the definite determiner *the*. The complement content role was typically an adjective (e.g., *brown*), but for a subset of items it consisted of a noun or noun phrase that served as the complement of the sentence predicate (e.g., *headphones* in *wore headphones*). The sentence set was comprised of 30 unique combinations of these four content roles with each corresponding to the same set of four pictures. We generated the specific sentences using custom MATLAB (MathWorks) code that constructed the sentence wording from the four content roles and the desired sentence structure. The final sentence set included 57 OR sentences, 45 SR sentences, 33 SMC sentences, and 25 OMC sentences. The number of trials per sentence structure slightly favored OR sentences due to scientific interest. All presented sentences belonged to one of these four experimental structure types. Overall the stimulus set was designed to maximize the number of informative trials while maintaining participant engagement and comprehension accuracy. The 30 unique content role combinations were presented in cycles where each one of the 30 content role combinations was presented in that cycle before starting the next cycle of combinations. We ensured that no instances of a content role combinations occurred within 10 trials of each other, which otherwise could occur at the cycle boundaries.

To induce variation in the verb tenses and the sentence and phrase lengths, we generated the sentences in four different verb tense/aspect combinations: simple present (e.g., “chases”), present progressive (“is chasing”), simple past (“chased”), and past progressive (“was chasing”). Each tense could be represented using the same set of pictures. The tense was randomly applied to each sentence independently of all other experimental manipulations. Each tense was used the same number of times in the sentences. In the OR and SR sentence types, the copular verb introducing the predicate (i.e., *is* or *was*) matched the tense of the main verb: *is* was used with the present-tense conditions, and *was* was used with the past-tense conditions.

To create variation in the part of speech present at the phrase boundary of OR sentences, we used prepositional verbs (e.g., *to speak to*) as the main verb in 33 of the 160 sentences, with the remaining main verbs being non-prepositional (e.g., *chases*).

To investigate how semantic information affects the processing occurring at phrase boundaries, we manipulated the semantic congruity of two aspects of the sentences. We manipulated the agent-patient mapping of the sentences to have 3 levels of semantic congruity: congruent (e.g. *The cop arrested the robber.*), reversible (*The lawyer that spoke to the judge was tall.*) and incongruent (*The robber arrested the cop.*). Similarly, we independently manipulated the complement-noun assignment of the sentences to have 3 levels of semantic congruity: congruent (e.g. *The striped tiger chased the lion.*), reversible (*The brown dog chased the cat.*) and incongruent (*The striped lion chased the tiger.*).

### Analyses

#### Preprocessing

We analyzed the per-word reaction times (RTs) reading the sentence as the primary dependent measure of interest. We also analyzed the picture choice of the participants at the end of the sentence and their response time making that picture choice as secondary dependent measures of interest. We excluded implausibly long RTs (>30 seconds) and the first-words of trials when the RT of the first word was unusually long (>0.7 seconds). The first word of each sentence always started with the word *the*, so latencies that long likely reflect task initiation and distraction at the onset of the sentence rather than sentence-level processing. Some OMC sentences became structurally ambiguous when constructed from items whose predicate contained a noun complement (e.g., *The cop arrested the robber wearing blue*). These trials were excluded from analyses involving the OMC condition (n=5).

For the across-participant analyses of Figure 5, for plotting purposes we threw out one outlier subject that had a baseline RT that was just over twice that of the next nearest participant. However, this participant was included in the associated statistical tests that we report, and their inclusion or exclusion did not affect the significance pattern of the results.

#### Mixed-effects regression overview

We analyzed the per-word RTs using linear mixed-effects models with random intercepts for participants and items that we fit using the fitglme function in MATLAB. We fit the models assuming a normal error distribution after log-transforming reading times, which provided a reasonable approximation to normality. We defined item-level random effects over the 30 unique semantic item combinations rather than the 160 individual sentence realizations of these because multiple sentence realizations shared the same underlying lexical-semantic content, allowing repeated observations of each item. The appropriate nuisance variables (described below) were included only as fixed effects in every model. When experimental predictors were included, we included them as both fixed effects and random slopes across participants and items. Sentence structure type (OR, SR, SMC, and OMC) was included as an experimental variable in the base model for all analyses. Additional candidate experimental predictors included word position, a phrase boundary indicator, the number of open nodes, the number of nodes closing, and the agent-patient and complement-noun semantic congruence reveal-point variables (described below).

Whenever an additional experimental predictor was included, its two-way interaction with sentence structure type was always also included as a fixed effect, reflecting the possibility that the predictor’s effect could differ across sentence structures. Corresponding random slopes for both the main effect and the interaction were included across participants. For items, random slopes were included only for the main effects of all the experimental predictors including sentence structure type and not for their interactions with sentence structure type. This more parsimonious random-effects structure was adopted because each semantic item combination was observed only five to six times, providing limited information to estimate item-specific interaction slopes reliably. Including interaction slopes across items would have substantially increased model complexity without sufficient repeated observations per item to support stable estimation.

We also explored models including additional three-way interactions of scientific interest; however, all such models failed to converge.

#### Semantic congruence reveal points

The semantic congruence variables were coded as four-level categorical predictors with the values *Not revealed yet*, *Congruent*, *Reversible*, and *Incongruent*. At the beginning of each sentence, both semantic congruence variables were coded as *Not revealed yet*. Once sufficient linguistic information had been presented to determine the semantic relationship of interest, the variable transitioned to the appropriate final state for the sentence (*Congruent*, *Reversible*, or *Incongruent*) and remained at that value for each word for the remainder of the sentence.

Because the point at which semantic congruence becomes available is not uniquely defined, we evaluated several plausible operational definitions of the reveal point for both the agent-patient and complement-noun congruence variables.

For the agent-patient congruence variable, we considered four reveal-point definitions:

1. At the main verb for all sentence types.
2. At the preposition following the main verb for prepositional-verb sentences, and at the main verb for all other sentence types.
3. After the first noun, second noun, and main verb had all been presented.
4. As in Option 3, but additionally requiring presentation of the preposition in prepositional-verb sentences.

For the complement-noun congruence variable, we considered two reveal-points:

1. After both the first noun and the complement had been presented.
2. After the first noun, second noun, and complement had all been presented.

#### Phrase boundaries

Each sentence structure has precisely one intra-sentence phrase boundary, which is the boundary marking the end of the subject noun phrase of the sentence. The word at this boundary, which is the last word of the subject noun phrase, is the first noun for SMC and OMC sentences, and the last word of the relative clause for OR and SR sentences. We considered the following 3 variations for the phrase boundary indicator variable in our models:

1. One indicator variable set to 1 at any phrase boundary during the sentence and at the sentence-final word.
2. Two separate indicator variables-one for the phrase boundary in the middle of the sentence and one for the sentence-final word.
3. Only an indicator variable for the major phrase boundary in the middle of the sentence.

#### Mixed-effects regression model-building procedure

During the initial stages of model development, we first determined the operational definitions of the semantic congruence reveal points and phrase boundary indicator variables. Because these variables are not uniquely defined, we evaluated the plausible definitions described above using our initial approximation of the final mixed-effects model. To determine the semantic congruence reveal-point definitions, we jointly evaluated all combinations of the agent-patient and complement-noun reveal-point definitions using a model that did not include a phrase boundary predictor. We then selected the combination yielding the lowest Akaike Information Criterion (AIC) for subsequent model-building analyses. We then evaluated the alternative phrase boundary indicator variables using the same model structure. The single “any phrase boundary” indicator (Option 1), which treated both within-sentence and sentence-final phrase boundaries as a single predictor, provided the best and most parsimonious model fit and was therefore adopted for all subsequent analyses.

Using these operational definitions, we performed forward model building beginning with a base model that included sentence structure type. At each step, the addition of each remaining candidate experimental predictor to the existing base model was evaluated separately. For each candidate predictor being evaluated, we fit two models using MATLAB’s fitglme function. In both models, the candidate predictor was added to the random-effects structure. One model then also included the predictor as a fixed effect (and the associated interaction with sentence structure type), whereas the other omitted these fixed effects. We then compared the two models in each pair of models using MATLAB’s compare method, which performs a likelihood-ratio test of the added fixed effects. In this way both models being compared always shared the same random effects structure, and we never tested experimental variables as fixed effects without corresponding random slopes for participants and items (Barr et al., 2013). We then ranked candidate predictors according to the significance of this likelihood-ratio test, with Bonferroni correction applied for the number of predictors under consideration at each model-building step. As a secondary measure of model fit, we also compared AIC differences between the paired models and we found it to agree with the likelihood-ratio test results in every instance. We then added to the model the predictor providing the strongest evidence for improved model fit and repeated the procedure with the remaining candidate predictors until no additional predictor significantly improved model fit.

Following completion of the forward model-building procedure, we re-evaluated the semantic congruence reveal-point definitions using the finalized model structure and selected for the final reported analyses the reveal-point definitions that provided the best overall model fit with the finalized model. Because the final model did not include a phrase boundary predictor, we did not re-examine the phrase boundary definitions. This re-evaluation resulted in a different optimal definition for the agent-patient reveal point than had been used during model building, whereas the optimal complement-noun reveal-point definition remained unchanged. Consequently, agent-patient reveal-point Option 1 and complement-noun reveal-point Option 1 provided the best fits in the final model and were used in all reported analyses.

#### Nuisance variables-word frequency

We obtained word frequency values from the SUBTLEX-US corpus (Brysbaert & New, 2009), specifically the part-of-speech annotated version of the database (Brysbaert et al., 2012). We derived part-of-speech-specific word frequencies from the reported frequencies for each lexical category. We used the base-10 logarithm of these frequency values in all analyses.

#### Nuisance variables-Residualized word length

As measures of word length, we considered both the number of letters and the number of syllables in each word. We ultimately used the number of letters because it correlated more strongly with participant reading times and is the more natural measure of length for visually presented text. Because word length and word frequency were strongly correlated, we residualized word length with respect to log word frequency. Specifically, we fit a linear regression predicting the number of letters from log word frequency across the unique words in the stimulus set and used the resulting residuals as the measure of word length in all subsequent analyses.

#### Nuisance variables-Surprisal

We computed word-level surprisal values using the distilgpt2 transformer language model (Radford et al., 2019; Sanh et al., 2019) as implemented in Hugging Face’s *transformers* library (v4.44.2). Experimental sentences were tokenized using the model’s byte-pair encoding vocabulary, and word-level surprisal was calculated as the sum of the negative log₂ probabilities (bits) of the constituent subword tokens.

We initially evaluated surprisal as a potential nuisance covariate because of its well-established effects on reading time. However, surprisal was strongly correlated with lexical frequency in the present stimulus set (Spearman’s ρ = −0.77, *p* < 10⁻²⁴⁴). Simultaneous inclusion of surprisal and lexical frequency produced unstable parameter estimates, consistent with substantial multicollinearity. We therefore also evaluated a residualized surprisal measure, obtained by removing variance attributable to lexical frequency, but this likewise yielded unstable parameter estimates. Given this substantial overlap, lexical frequency was retained as the lexical control variable because it produced stable parameter estimates across models, and surprisal was omitted from the final statistical analyses.

#### Nuisance variables-Baseline RT

In order to get a measure of the baseline state of the participant going into each trial, we defined for each sentence structure a baseline period over the first few words of the sentence before any experimentally consequential words in the sentence were reached, with different definitions of the baseline period for each of the sentence structure types. For OR sentences, we defined this as everything before the second noun in the sentence (i.e., everything from *The* up to and including *the* in: *The dog that the cat chased was brown*.). For SR, SMC and OMC sentences, we defined this as everything before the main verb in the relative clause (i.e., everything from *The* up to and including the first *was* in: *The dog that was chasing the cat was brown.*, or in *The brown cat was chasing the dog*, or in *The cat was chasing the brown dog.*).

#### Analysis conventions

Unless otherwise indicated, figures display baseline-subtracted word-RT residuals to better visualize the effects of interest. To generate these plots, we first computed word-RT residuals using the fitglme function and its residuals method after accounting for the fixed effects of log word frequency and residualized word length. For each trial, a baseline residual was then calculated as the mean residual over the sentence baseline period and subtracted from the residual of every word in that trial. This baseline subtraction reduces the influence of trial-to-trial fluctuations in overall response speed while preserving within-sentence effects.

All statistical analyses of word RT were performed on natural log-transformed word RTs, which yielded approximately normally distributed data appropriate for the statistical tests we employed. Neither residualization nor baseline subtraction was applied to the data used for statistical inference. Instead, the corresponding sources of variance were accounted for directly within the mixed-effects models through the inclusion of the appropriate nuisance variables.

#### Question Accuracy and RT analyses

We analyzed overall comprehension accuracy using a generalized linear mixed-effects model with a binomial error distribution. The model included random intercepts for participants and items (defined as in the word RT analyses) and sentence structure type as a fixed effect, with random slopes for sentence structure type across both participants and items. To compare the distribution of comprehension error types across sentence structure types, we fit a multinomial regression model among the error responses using the fitmnr function in MATLAB with effects coding. Complement-error served as the reference error category.

## Results

### Question Accuracy and RT

A generalized linear mixed-effects model revealed a significant effect of sentence structure on comprehension accuracy (**Figure 2a.,** F(3,5421) = 27.25, p < 0.001). As anticipated participants committed the most errors on OR sentences, although they broadly still showed a reasonably high comprehension rate for those sentences (89.4%). The distribution of error types significantly differed across sentence structure types (χ²(6) = 28.7. p<0.001). The proportion of “Both” error types (where participants selected the picture that was incorrect in terms of both agent-patient mapping and complement-noun mapping) relative to the complement error reference type was significantly elevated in OR sentences (Wald *z* = 2.50, *p* = .012). SMC sentences also yielded significantly lower relative proportions of “Both” errors (Wald *z* = 2.45, *p* = .014) and agent-patient errors (Wald *z* = 2.45, *p* = .014) relative to the complement error type. When comparing later epochs in the experimental session to earlier epochs (**Figure S1**), although accuracy improve in later relative to earlier epochs, the relatively decreased accuracy for OR sentences remained, and the tendency to commit “Both” errors when committing a comprehension error in OR sentences generally remained as well.

**Figure 2.**
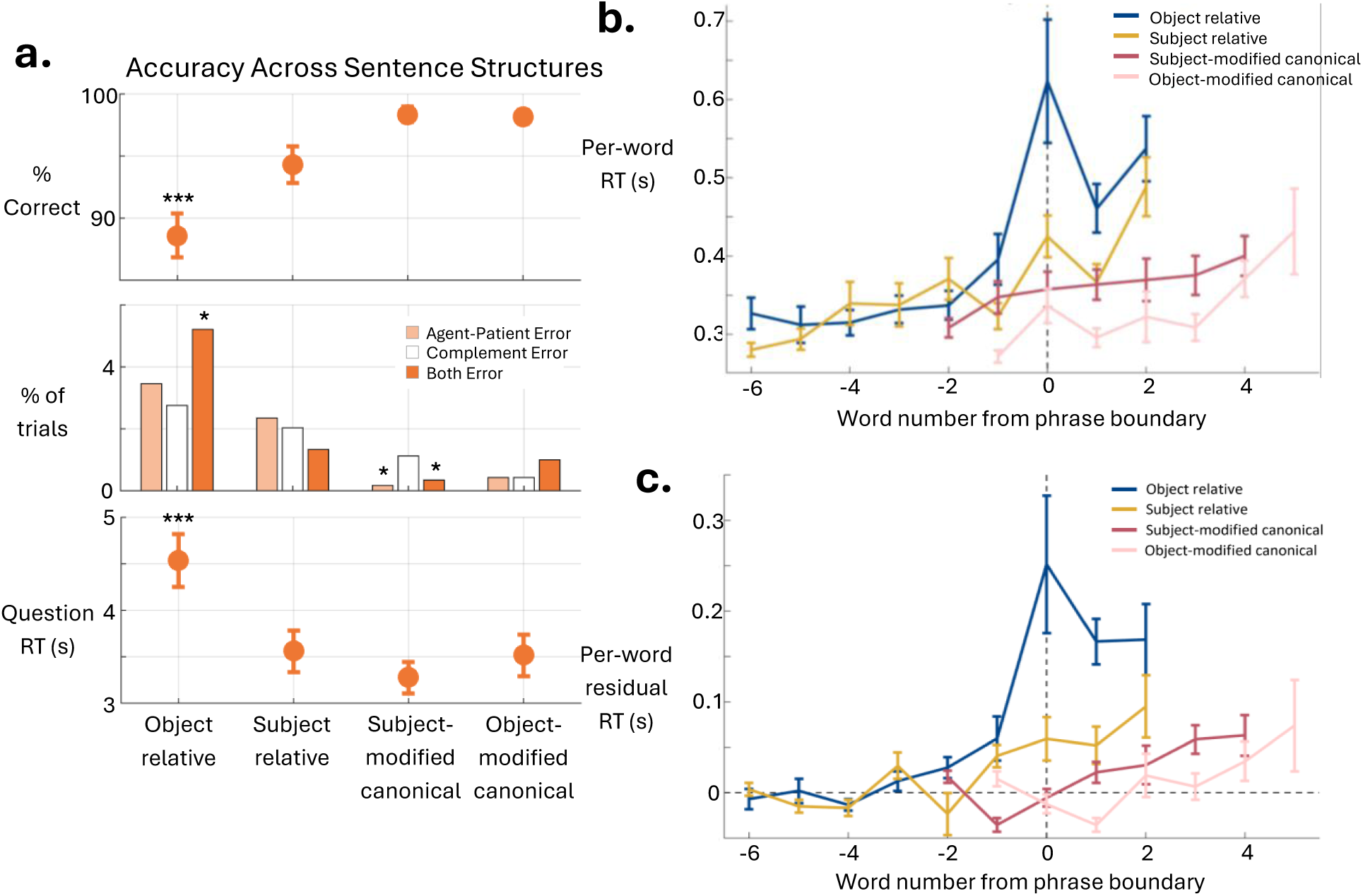
**(a)** The topmost graph displays the percentage participants responded correctly to across sentence structure types. The middle graph displays the percentage of trails that displayed agent-patient error, complement error, or both errors together. The bottommost graph displays the response times to the comprehension picture questions across sentence types. **(b)** The graph shows response time per-word for the four sentence structures, plotted by the word number from the phrase boundary. **(c)** The graph shows residual response time per-word for the four sentence structures after accounting for the effects of word frequency and word length with baseline RT subtraction, plotted by the word number from the phrase boundary.

### Per-Word RT

We aligned the per-word RTs at the major phrase boundary of each sentence, which was the last word of the relative clause for OR and SR sentences and the first noun for SMC and OMC sentences, for raw as well as baseline-corrected residual per-word RTs (**Figure 2b-c**.). Participants show considerable RT increases at the phrase boundary specific to OR sentences, and to a lesser extent, SR sentences. RTs after the phrase boundary remain elevated for these sentence structures. The phrase boundary peak for OR sentences persisted through-out the session even while participants’ RT globally decreased in later epochs (**Figure S2**).

OR sentences typically have verbs at the phrase boundary (e.g. *chased* in *The dog that the cat chased was brown.*). However OR sentences with prepositional verbs have a preposition at the phrase boundary (e.g. *to* in *The woman that the waiter spoke to was short.*). We thus compared the phrase-boundary-aligned word RTs between prepositional verb and non-prepositional verb sentences to observe if the effect at OR phrase boundaries was due to the verb or due to the phrase boundary. The results in **Figure 3** show RTs at the phrase boundary preposition elevated above the RT of the main verb in OR sentences, showing that the maximal increase in reading time is aligned with the phrase boundary rather than the verb itself.

**Figure 3.**
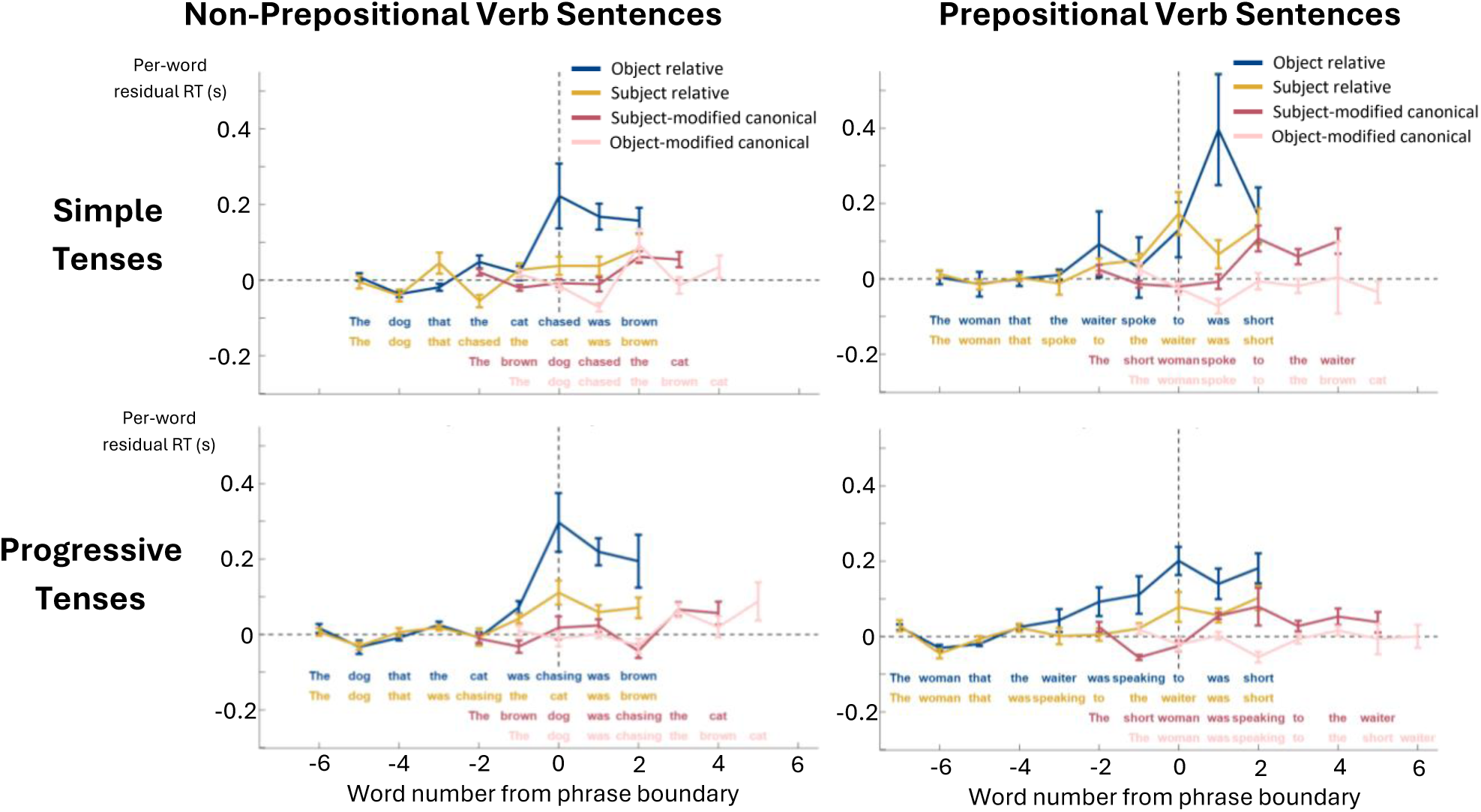
The four graphs display the residual response times per-word for each of the four sentence structures, placed by the word number from the phrase boundary. Sample sentences from the experiment are placed along to show this comparison clearly. The upper-left graph shows results for non-prepositional verb sentences with simple tenses. The lower-left shows results for non-prepositional verb sentences with progressive tenses. The upper-right shows results for prepositional verb sentences with simple tenses. The lower-right shows results for prepositional verbs with progressive tenses.

As another way to inspect these effects, for each of the sentence structure types we grouped the word RTs according to their semantic content role within the sentence (the first noun, the verb, the second noun, and the noun-complement), with comparative categories added for function words (e.g., determiners and auxiliary verbs) and the preposition following the verb in prepositional verb sentences (**Figure 4**). For this analysis, we excluded words immediately following phrase boundaries in relative-clause sentences to minimize contamination of the function-word category by spillover from phrase-boundary processing. We ran two mixed-effects regression models including sentence structure type, content word type, and their interaction as predictors of interest. Separate models were fit for non-prepositional verb sentences and prepositional verb sentences. The results revealed a significant interaction and main effects for both content word type and structure type for both non-prepositional verb sentences (interaction: F(12, 29192) = 6.62; p < 0.001, content word type: F(4, 29192) = 6.34; p<0.001, structure type: F(3, 29192) = 5.70, p < 0.001) and prepositional verb sentences (interaction: F(16, 8617) = 6.62; p < 0.001, content word type: F(4, 8617) = 6.34; p<0.001, structure type: F(3, 8617) = 5.70, p < 0.001). To characterize the significant interaction between sentence structure and content word type, we performed simple-effects tests of sentence structure within each content-word type using model contrasts separately for non-prepositional verb sentences and prepositional verb sentences. Significant simple effects were followed by pairwise comparisons between sentence structure types. These results are summarized in **Table 1**. For non-prepositional verb sentences, the most prominent effect was the increased RT at the verbs of OR sentences, where the phrase boundary of those sentences occurs. Notably, for prepositional verb sentences the most prominent effect was the increased RT at the prepositions of OR sentences, where the phrase boundary of those sentences occurs, although the RT of verbs in these sentences was also higher than in other sentence structures. This pattern suggests that most of this added processing underlying the object-relative pause observed near the phrase boundary occurs when the phrase itself is completed, with some of the processing occurring when the verb happens even before the phrase is completed. The significant increased RT at the complement for both relative clause sentence structure types relative to the canonical sentences may reflect additional confusion in the complement-to-noun assignment for these sentences as well as generally elevated RTs that seem to be present for the rest of the sentence after the phrase boundary for these sentences (**Figure 2**). The significantly increased RT at N2 for SR sentences relative to OR sentences corresponds to the phrase boundary that occurs for SR but not OR sentences at that word. The increased RT at the verb for SMC relative to OMC sentences may reflect additional working memory load relative to the additional complement that has already been received in the SMC sentences.

**Figure 4.**
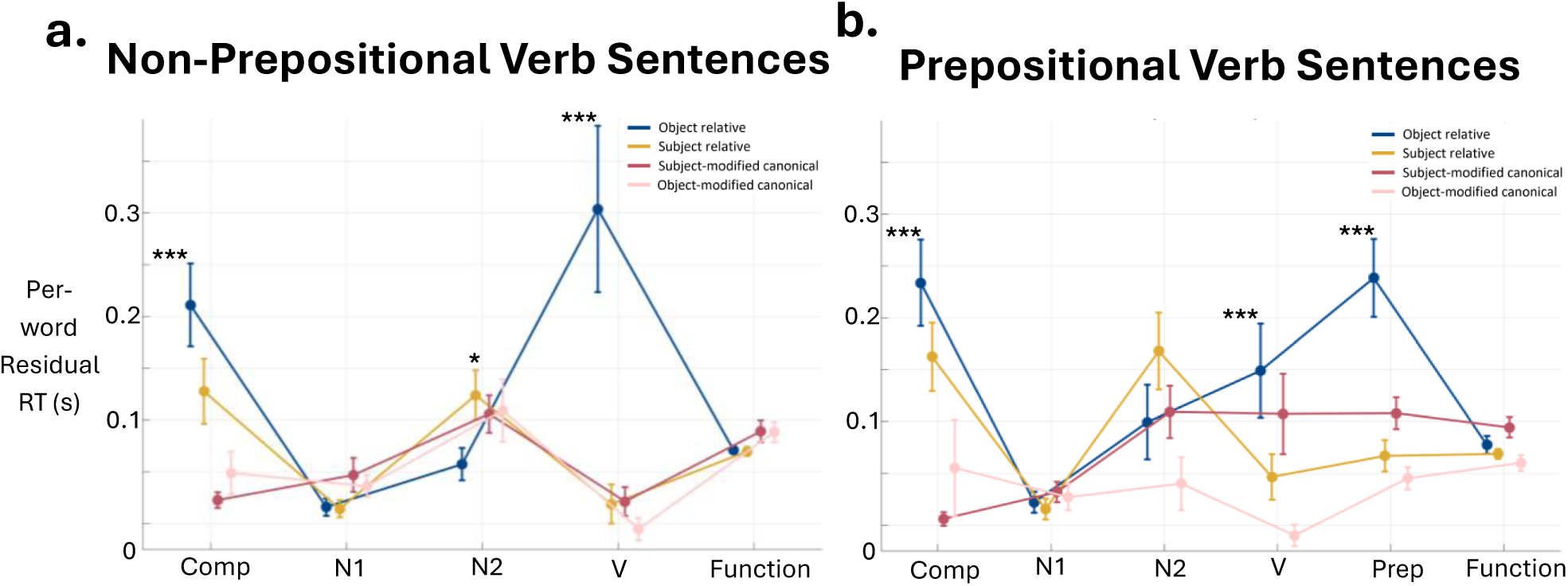
The two graphs show the residual response times across parts of speech within the four sentence structures. **(a)** The graph displays that of non-prepositional verb sentences, including the following parts of speech: complementizer, first noun, second noun, verb, and following functional word. **(b)** The graph displays that of prepositional verb sentences, including the same parts of speech but adding the preposition. Words immediately following phrase boundaries in relative-clause sentences were excluded from this analysis.

**Table 1.**

| Content word type | Simple-effect test of sentence structure | Significant pairwise comparison | F | p |
| --- | --- | --- | --- | --- |
| **Prepositional Verb Sentences** |  |  |  |  |
| Complement | $F(3, 8617)=19.38; p<0.001$ | OR > SMC | 52.47 | <.001 |
|  |  | OR > OMC | 37.61 | <.001 |
|  |  | SR > SMC | 28.13 | <.001 |
|  |  | SR > OMC | 14.51 | <.001 |
| Verb | $F(3, 8617)=11.59; p<0.001$ | OR > SR | 6.76 | 0.009 |
|  |  | OR > SMC | 8.95 | 0.003 |
|  |  | OR > OMC | 33.62 | <.001 |
|  |  | SMC > OMC | 6.71 | 0.01 |
| Preposition | $F(3, 8617)=14.09; p<0.001$ | OR > SR | 13.84 | <.001 |
|  |  | OR > SMC | 18.14 | <.001 |
|  |  | OR > OMC | 41.3 | <.001 |
| Function | $F(3, 8617)=7.07; p<0.001$ | OR > OMC | 21.05 | <.001 |
| **Non-prepositional Verb Sentences** |  |  |  |  |
| Complement | $F(3, 29192) = 10.87; p < .001$ | OR > SR | 3.98 | 0.046 |
|  |  | OR > SMC | 23.5 | <.001 |
|  |  | OR > OMC | 26.6 | <.001 |
|  |  | SR > SMC | 20.36 | <.001 |
|  |  | SR > OMC | 20.43 | <.001 |
| N2 | $F(3, 29192) = 3.19; p = 0.023$ | SR > OR | 9.2 | 0.002 |
| Verb | $F(3, 29192) = 8.05; p < .001$ | OR > SR | 15.96 | <.001 |
|  |  | OR > SMC | 17.76 | <.001 |
|  |  | OR > OMC | 23.6 | <.001 |
|  |  | SR > OMC | 5.34 | 0.021 |

### RT Spillover Effect

We observed an apparent spillover effect in which, for some participants, the first word following the OR phrase boundary exhibited a larger RT increase than the phrase-boundary word itself. We hypothesized that this spillover effect would be more pronounced in participants with faster baseline reading speeds because these participants are more likely to initiate the button press for the next word before processing at the phrase boundary has fully completed. Under this account, processing triggered by the phrase boundary continues into the following word, shifting a portion of the RT cost from the phrase-boundary word onto the subsequent word. In **Figure 5a**. we plotted for each participant the mean RT on the words after the phrase boundary in OR sentences minus the RT at the phrase boundary word of these sentences. Across participants, larger baseline RTs were associated with smaller spillover effects at the first word following the phrase boundary (Spearman’s ρ = -0.53, *p* = .001). In contrast, baseline RT was unrelated to OR sentence comprehension accuracy (Spearman’s ρ = -0.18, p = .301) or overall comprehension accuracy (Spearman’s ρ = 0.12, p = .499), suggesting that the relationship between baseline reading speed and spillover is unlikely to simply reflect differences in task engagement or comprehension performance and is instead consistent with stable individual differences in how participants paced their responses during self-paced reading. **Figure 5b** and **5c** show the phrase-boundary-aligned word RT profile for the slow-baseline and fast-baseline participants using 0.29 seconds as a baseline RT cutoff.

**Figure 5.**
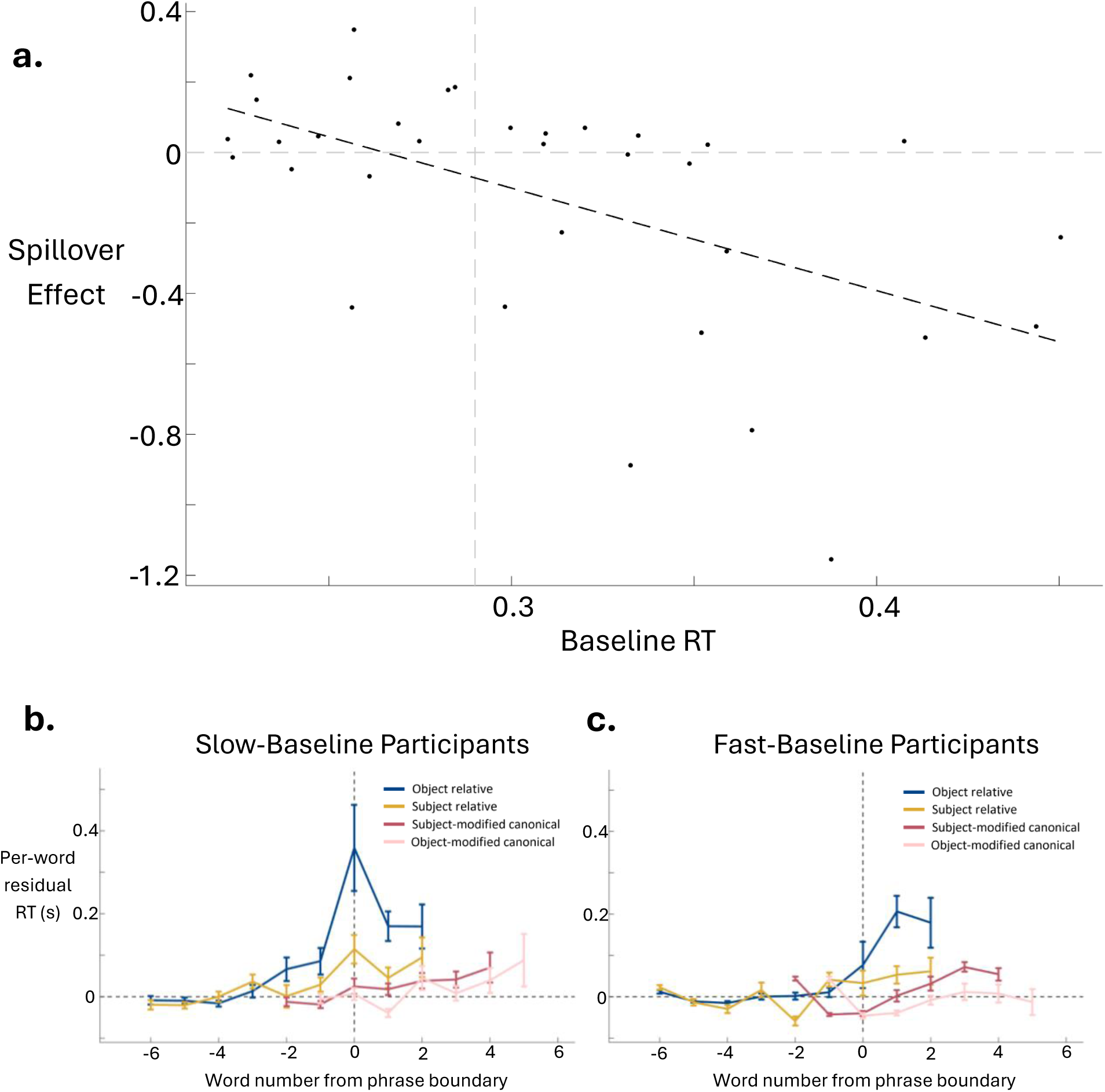
**(a)** The graph displays the spillover effect as a function of the baseline response times. **(b)** The graph shows the per-word residual response times for slow-baseline participants of four sentence structures plotted by the word number distance from the phrase boundary. **(c)** The graph shows the per-word residual response times for fast-participants of four sentence structures plotted by the word number distance from the phrase boundary.

### Node closing analyses

To evaluate evidence for the node-tracking framework in self-paced reading RTs, we analyzed the effects of the number of closing nodes on relative-clause sentences at the phrase boundary at the end of the subject noun phrase, at the word after the phrase boundary (based on the number of nodes closing at the phrase boundary immediately preceding it), and at the sentence-final word (**Figure 6**). We tested this using mixed-effects linear models with the parameter of the number of nodes closing as a fixed effects parameter with random slopes across participants and items applied to each of these words across the dataset. There was a significant positive effect at the phrase boundary word (t(3376) = 3.78; p<.001), but the effect was not significant for the word after the phrase boundary (p=.857) or the sentence-final word (p=.920).

**Figure 6.**
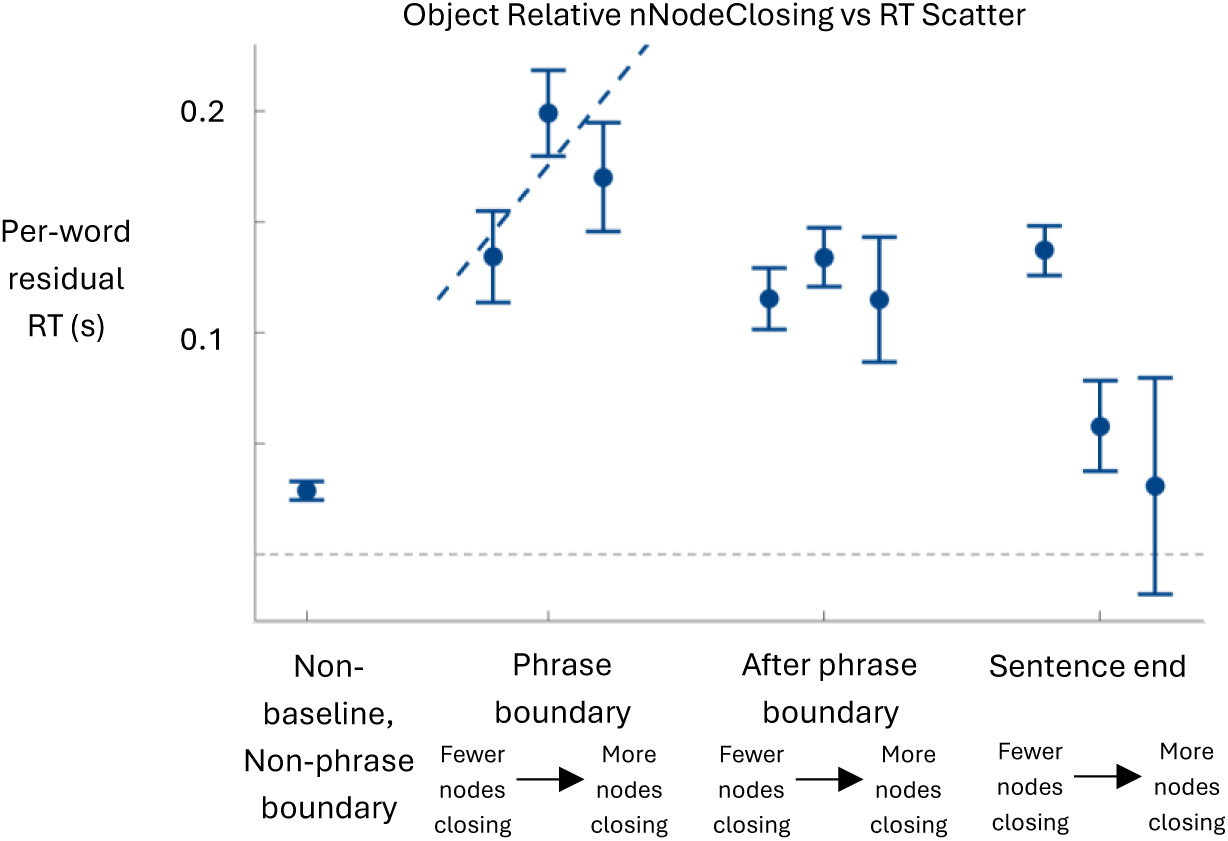
The graph displays the per-word residual response times for each syntactic region for the object relative sentences. Excluding the non-baseline non-phrase boundary word, which is shown as a reference point, each syntactic region includes three data points ordered to show fewer to more nodes closing.

### Canonical sentence analyses

SMC and OMC sentences differ in the position of the complement and the noun they modify-for SMC sentences the complement occurs before the main verb and modifies N1 while in OMC sentences this occurs after the verb and modifies N2. We compared the two to see if these differences result in subtle word RT difference between the two structures. We compared the RTs at N1 and N2 in separate analyses, restricted to instances when the complement was an adjective. Thus in each case this amounts to comparing, e.g. *the brown dog* to *the dog* in instances where that noun phrase is the subject noun phrase at the beginning of the sentence in one analysis and the object noun phrase at the end of the sentence in another analysis. The results are shown in **Figure 7a** and **7b**. Using a mixed effects model applied to these data with structure type as the only experimental predictor, the difference was not significant for the first noun phrase (p=0.436) or the second noun phrase (p=0.944). We further tested for a linear increase in RT across the *Det-Adj-N* triplets pooled main between the SMC and OMC sentences, using linear mixed effects model with Phrase Position within this triplet (a variable that is 1 for the Determiner, 2 for the adjective and 3 for the noun) as the only experimental predictor. The effect was positive and significant (F(1,4234) = 6.9994; p = .008). This effect is seen more clearly when comparing the raw word RT data between these conditions (**Figure S3**).

**Figure 7.**
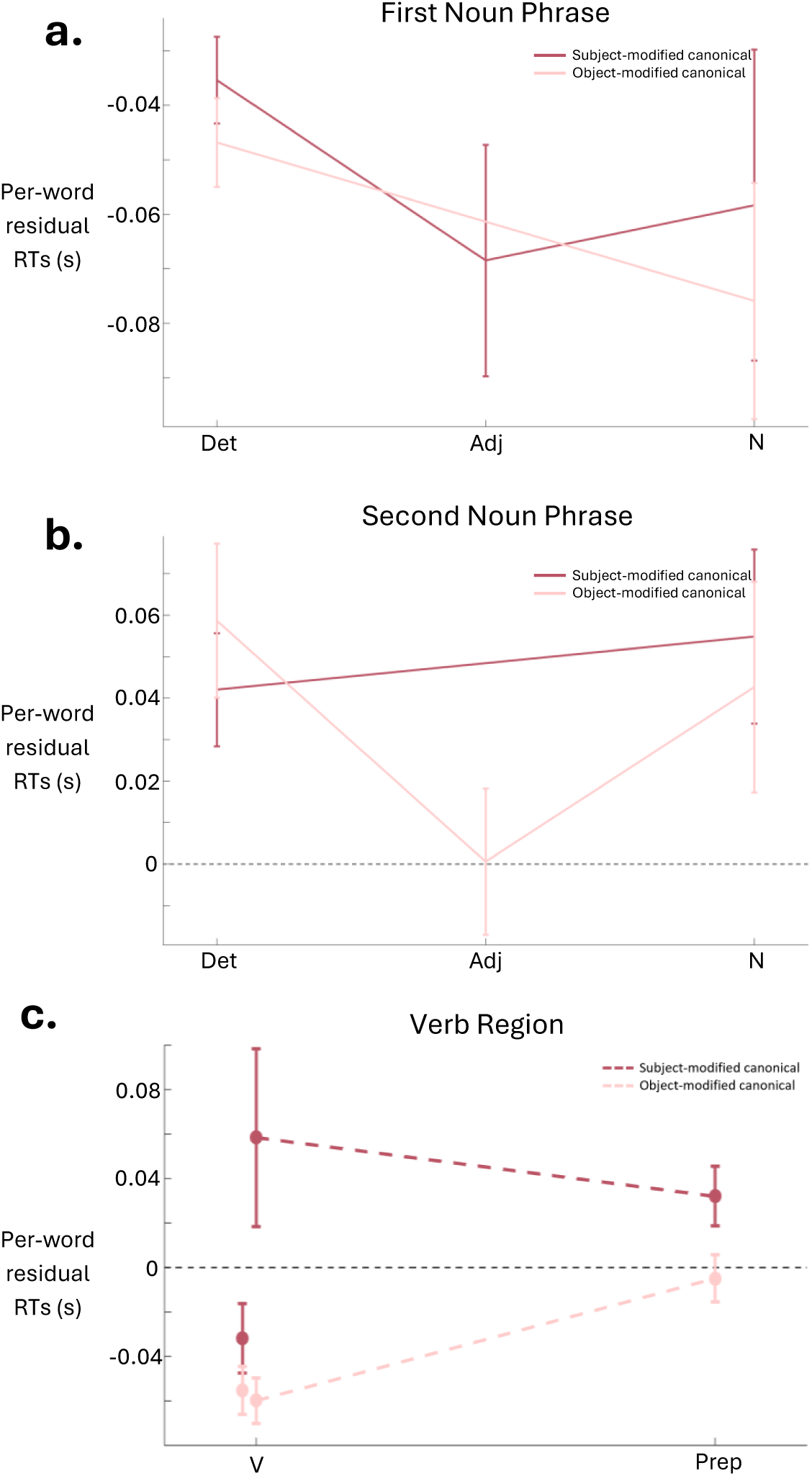
The three graphs describe the per-word residual response times for subject-modified canonical and object-modified canonical sentences. **(a)** The graph describes the first noun phrase in the sentence structures, including the phrase’s determiner, the adjective, and the noun. The graph does not involve baseline subtraction because the baseline for these sentences would include this noun phrase. Baseline subtraction was also not included as a factor in the corresponding mixed effects model used for significance testing this analysis. **(b)** The graph describes the second noun phrase in the sentence structures, including the phrase’s determiner, the adjective, and the noun. **(c)** The graph describes the verb region in the sentence structures, including the verb and following preposition.

We also plotted the effects for the verb region, including the verb for non-prepositional verb sentences and the main verb plus the preposition for prepositional verb sentences (**Figure 7c**). Using a mixed effects model applied to these data with structure type as the only experimental predictor, the result approached significance, with SMC sentences having larger RTs in the verb regions than OMC sentences, but the effects was not ultimately significant (p=0.077).

### Semantic congruence analyses and regression model-building analysis

Semantic congruence of the agent-patient mapping had a large apparent effect for object relative sentences in particular (**Figure 8**), being best fit by an onset of the effect at the verb that persists from there through the rets of the sentence. The effect follows the anticipated pattern with the incongruent trials showing the longest RTs, the congruent trials with the shortest RTs and the reversible trials with

**Figure 8.**
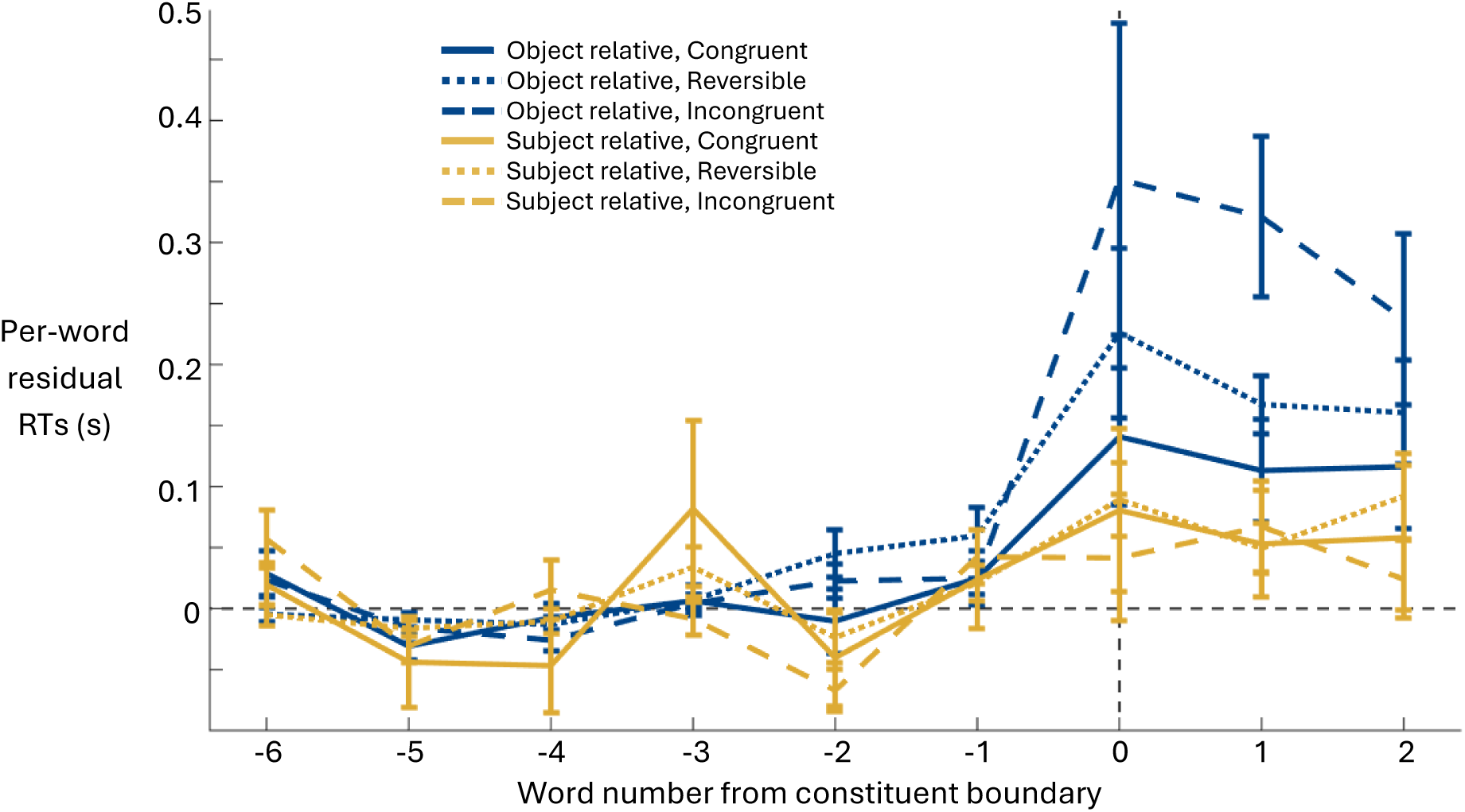
The graph displays the residual response times per-word plotted by their word number from the constituent boundary. Described are the object relative and subject relative sentence structures and their plausibility (congruent, reversible, incongruent).

RTs in the middle. For completeness in **Figure S4** we show the results for the two canonical sentence structures grouped according to the semantic congruence of the agent-patient mapping, for which there was no effect.

The final model resulting from the model building procedure we employed is shown in **Table 2**. The semantic congruence of the agent-patient mapping was the first experimental term added to the model, followed by the semantic congruence of the noun-complement assignment, and finally followed by the numbers of closing nodes, after which no further experimental parameters exceeded the significance threshold when considered.

**Table 2.** Analysis of variance for the final linear mixed-effects model. Only experimental predictors are shown. The model additionally included log word frequency, residualized word length, baseline RT, and part of speech as nuisance covariates, together with the random effects described in the Methods.

| Predictor | df | F | p |
| --- | --- | --- | --- |
| Structure Type | 3, 39006 | 6.91 | $1.20 \times 10^{-4}$ |
| Number of Nodes Closing | 1, 39006 | 0.02 | 0.894 |
| Patient–Agent Semantic Congruence Revealed | 3, 39006 | 4.68 | 0.00285 |
| Noun-Complement Semantic Congruence Revealed | 3, 39006 | 2.62 | 0.0487 |
| Structure Type × Number of Nodes Closing | 3, 39006 | 3.39 | 0.0173 |
| Structure Type × Patient–Agent Semantic Congruence Revealed | 9, 39006 | 9.87 | $2.93 \times 10^{-15}$ |
| Structure Type × Noun-Complement Semantic Congruence Revealed | 9, 39006 | 4.05 | $3.33 \times 10^{-5}$ |

## Discussion

In this study, we used self-paced reading to investigate how syntactic structure, semantic interpretation, and phrase structure jointly influence online sentence processing. Across several complementary analyses, object-relative (OR) sentences consistently produced the greatest processing difficulty, reflected by increased reading times at the completion of the relative clause, reduced comprehension accuracy, and an increased tendency to commit errors involving both agent-patient assignment and complement assignment simultaneously. These findings replicate the well-established processing cost associated with object-relative clauses while providing a more detailed characterization of when during sentence comprehension this cost emerges.

One of the clearest findings was that the largest increase in reading time occurred specifically at the completion of the subject noun phrase rather than being tied to a particular lexical category. OR sentences with non-prepositional verbs showed their largest RT increase at the main verb, whereas OR sentences with prepositional verbs instead showed the largest increase at the phrase-final preposition. Because these two sentence types differ only in the identity of the phrase-final word, this finding strongly suggests that the critical processing is triggered by completion of the phrase rather than by processing of the verb itself. Some increased processing was already apparent at the main verb of prepositional-verb sentences, suggesting that processing may begin as soon as sufficient syntactic information becomes available but reaches its maximum when the phrase is completed.

Our node-closing analysis further supports this interpretation. At phrase boundaries, reading times increased as a function of the number of syntactic constituents completed at that word, whereas no such relationship was observed at the following word or at sentence-final boundaries. This selective effect is consistent with the proposal that completion of syntactic constituents incurs an incremental processing cost during reading. Together with the phrase-boundary analyses, these results suggest that self-paced reading captures ongoing syntactic integration processes rather than simply reflecting lexical access or generalized sentence difficulty.

An additional observation concerns the substantial spillover of the phrase-boundary effect into the following word for many participants. Participants with faster baseline reading speeds exhibited significantly greater spillover despite showing no reduction in comprehension accuracy. This finding is consistent with the idea that faster readers often initiate the button press for the following word before processing associated with the phrase boundary has fully completed. Consequently, some of the processing cost becomes expressed on the subsequent word rather than the phrase-boundary word itself. This observation illustrates an important characteristic of self-paced reading data: the temporal location of processing costs may depend partly on individual response strategies, even when the underlying linguistic computation is similar.

One of the most interesting findings of the present study was the interaction between semantic congruence and syntactic structure. Agent-patient semantic congruence produced its largest effects specifically for object-relative sentences, whereas the effect was considerably smaller for the other sentence structures. This pattern suggests that semantic information does not simply contribute a constant processing cost whenever an implausible event is encountered. Instead, semantic interpretation appears to interact with the syntactic computations that are particularly demanding in object-relative clauses.

Object-relative sentences require readers to establish a non-canonical mapping between grammatical roles and semantic roles while simultaneously maintaining multiple noun phrases across the relative clause. Under these conditions, agent-patient semantic information may become especially important for resolving or confirming the intended interpretation. Plausible semantic relationships appear to facilitate this process, whereas semantically incongruent relationships increase the processing required at precisely the point where syntactic integration is most demanding. The relative absence of comparable semantic effects in subject-relative sentences argues against a purely lexical or semantic explanation and instead suggests that semantic information specifically modulates the computations involved in constructing the syntactic representation of object-relative clauses. More broadly, these findings provide an example of the close interaction between syntactic parsing and semantic interpretation during incremental sentence comprehension rather than these processes operating independently. Interestingly, the agent-patient congruence variable entered the regression model before any other experimental predictor during the forward model-building procedure, indicating that it explained more variance than either complement congruence or node-closing effects. This emphasizes that semantic interpretation contributes substantially to online processing even after controlling for lexical variables and sentence structure, while the strong interaction with OR sentences indicates that this contribution is closely linked to syntactic difficulty rather than representing a simple additive semantic effect.

In contrast, the complement-congruence effect should be interpreted more cautiously. Although statistically reliable in the regression analysis, its direction differed across sentence structures, with subject-relative sentences exhibiting the opposite pattern from that predicted. At present we do not have a satisfactory explanation for this finding. Consequently, we view the complement-congruence result as preliminary and believe it warrants replication before drawing strong theoretical conclusions.

The analyses comparing the two canonical sentence structures also provide several informative observations. Overall differences between subject-modifier canonical (SMC) and object-modifier canonical (OMC) sentences were relatively subtle, suggesting that these structures impose broadly similar processing demands. Nevertheless, Figure 4 demonstrated significantly increased reading times in the verb region for SMC sentences relative to OMC sentences in prepositional-verb items, consistent with the prediction that SMC sentences require maintenance of a completed modifier before processing the verb. Although this effect was modest and was not consistently detected across all analyses, its direction agrees with the hypothesis that the additional working-memory demands imposed by SMC sentences produce a measurable, albeit relatively small, processing cost.

The present findings have several methodological implications for studies using self-paced reading. First, analyses aligned to linguistically meaningful events, such as phrase boundaries, may reveal processing effects that are obscured when data are analyzed solely according to absolute word position. Second, because processing frequently spills over beyond the triggering word, particularly for faster readers, analyses that examine only a single word may underestimate the true magnitude of processing associated with a linguistic event. Finally, the observed sensitivity of self-paced reading to constituent completion and node-closing measures demonstrates that this relatively simple behavioral paradigm can provide surprisingly detailed information about the incremental computations underlying sentence comprehension. Several limitations should also be acknowledged. The sentence materials were intentionally designed to maximize experimental control and permit detailed manipulation of syntactic and semantic variables, which necessarily limits the range of sentence constructions examined. Likewise, the current analyses focus primarily on reading times and behavioral performance rather than directly measuring the neural mechanisms underlying these effects. Future studies combining comparable sentence manipulations with electrophysiological or intracranial recordings could help identify the neural processes responsible for the phrase-boundary effects observed here.

In summary, the present study demonstrates that self-paced reading is sensitive to the incremental processing associated with syntactic constituent completion, particularly during comprehension of object-relative clauses. The results indicate that processing costs are tied more closely to phrase completion than to individual lexical categories, scale with the amount of syntactic structure completed at a boundary, and interact strongly with agent-patient semantic interpretation specifically when syntactic demands are greatest. Together, these findings support a view of sentence comprehension in which syntactic structure building and semantic interpretation proceed incrementally and interact continuously throughout online language processing.

## Declaration of generative AI and AI-assisted technologies in the manuscript preparation process

During the preparation of this work, the authors used ChatGPT (OpenAI) to assist with language editing, improving the clarity of the manuscript, and provide feedback on aspects of the data analysis. After using this tool, the authors independently reviewed and evaluated the output, edited the content as needed, and take full responsibility for the analysis, interpretation, and content of the published article.

## Supplementary figures

**Figure S1.**
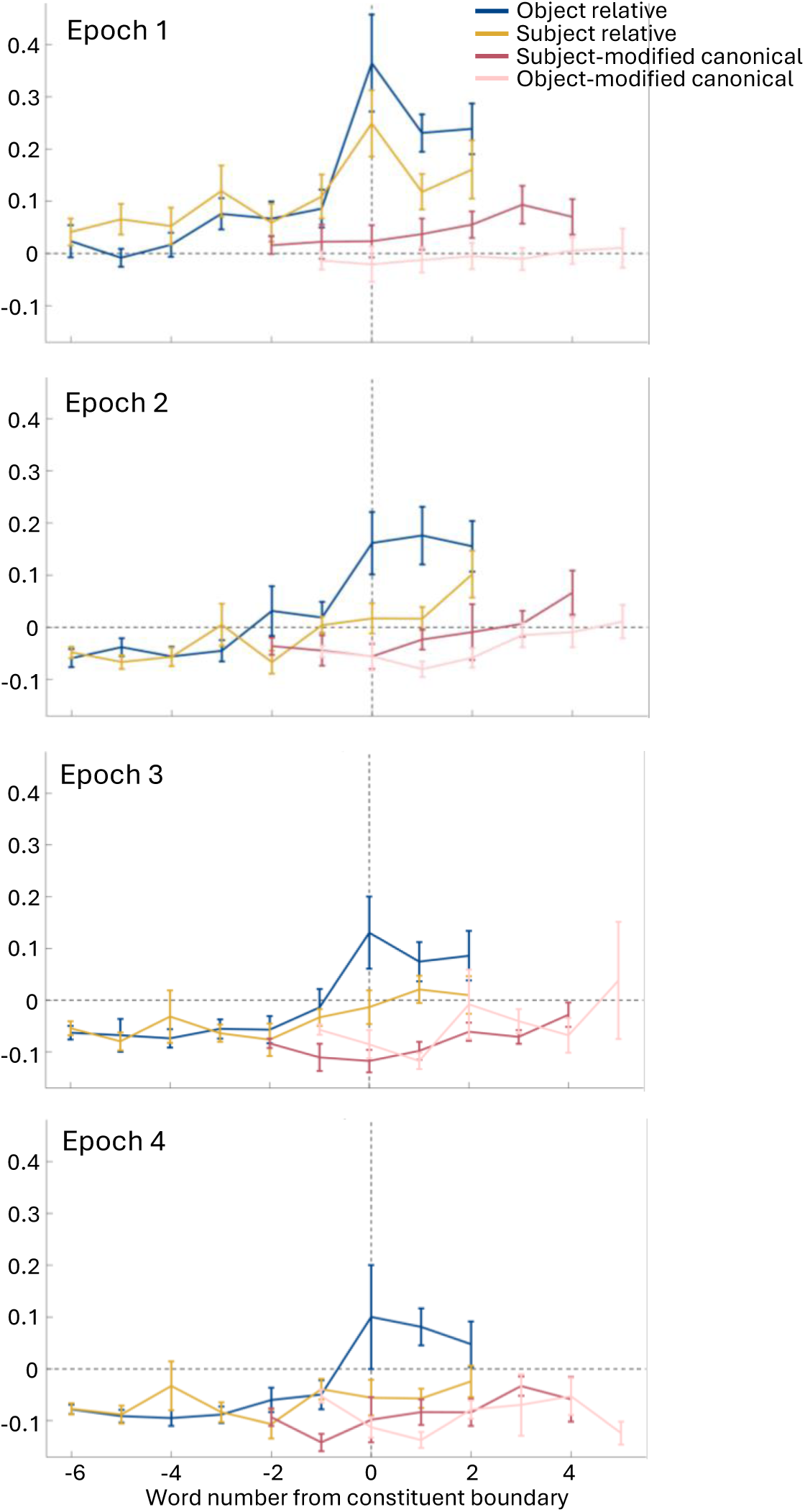
The graphs compare plots per-word response times across the word number from the constituent boundary to describe behavior over four succeeding epoch times.

**Figure S2.**
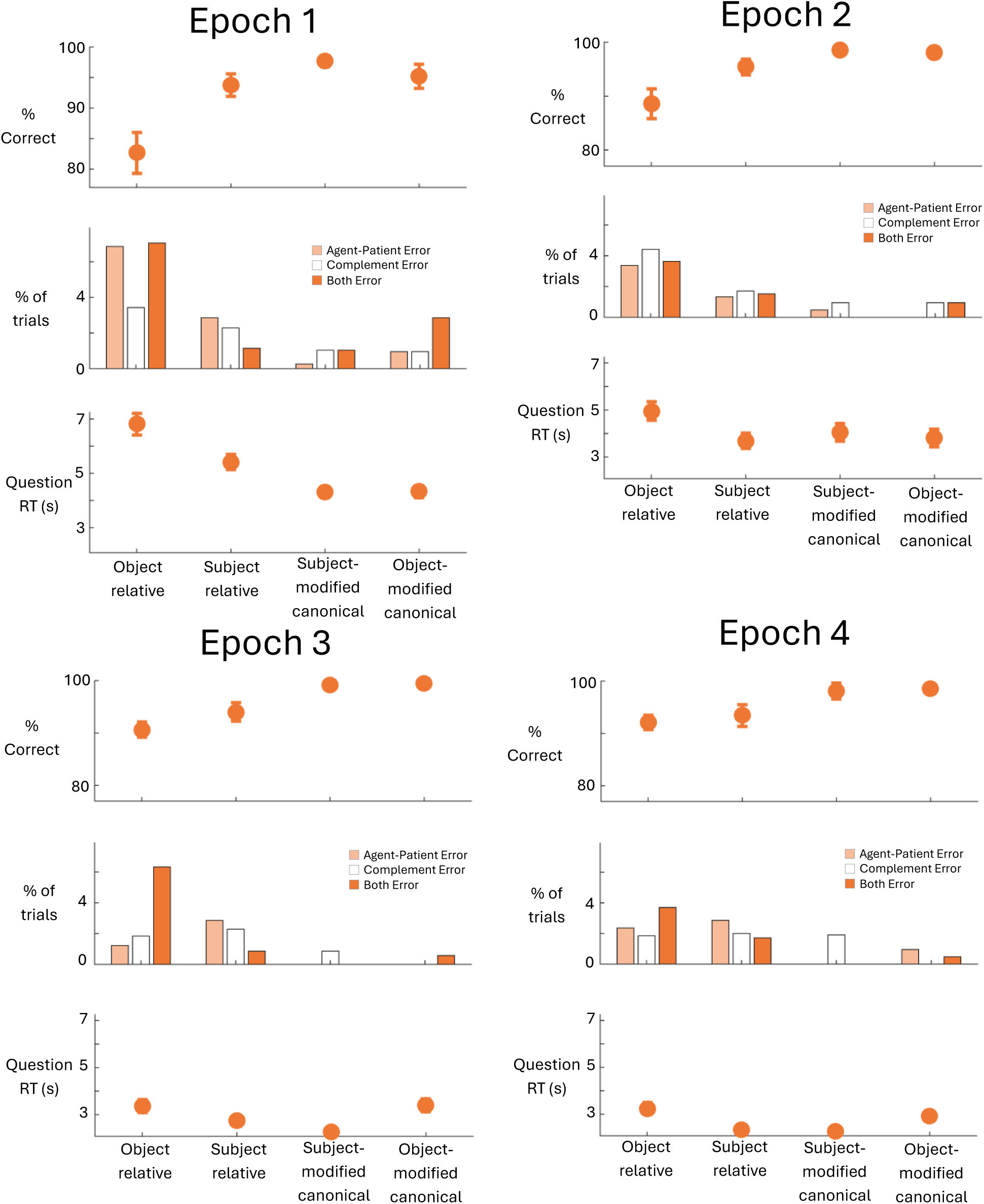
The graphs compare four different succeeding epochs across the four sentence structures: object relative, subject relative, subject-modified canonical, and object-modified canonical. Each epoch contains three graphs comparing behavior response to the comprehension questions. The topmost graph describes the percentage of accuracy for the comprehension questions. The middle graph describes the percent of trials that exhibited a agent-patient error or complement error, or both. The bottommost graph describes the response time of answering the comprehension questions.

**Figure S3.**
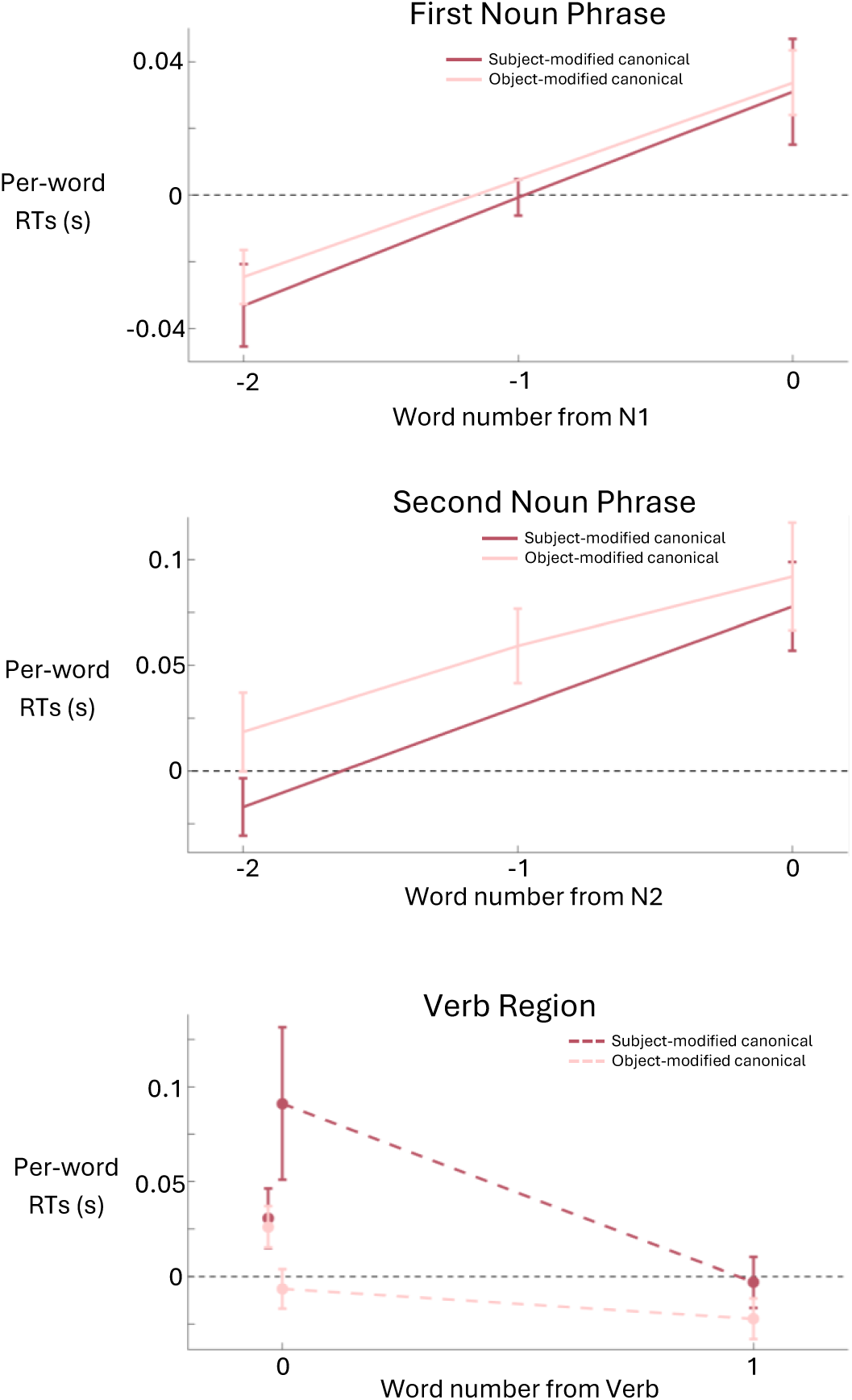
The three graphs describe the baseline-subtracted raw (not residualized) per-word response times for subject-modified canonical and object-modified canonical sentences. The top-left graph describes the RTs for the word number from the first noun phrase (N1). The bottom-left graph describes the RTs from the second noun phrase (N2). The top-right graph describes the word number from the verb.

**Figure S4.**
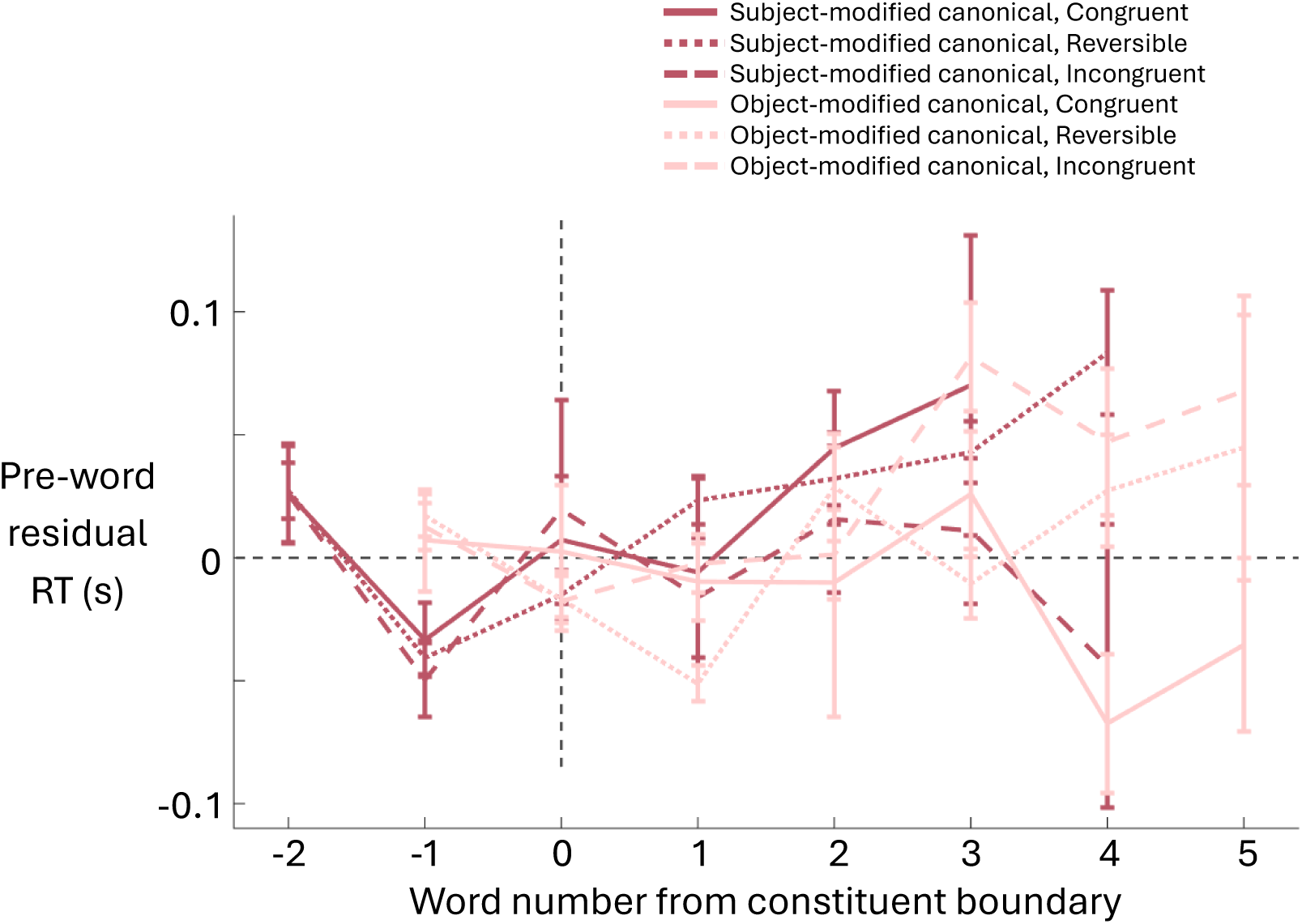
The graph describes the per-word residual response times by the word number distance from the constituent boundary for the subject-modified canonical and object-modified canonical sentence structures and their plausibility (congruent, reversible, incongruent).

